# A genome-wide association study for host resistance to Ostreid Herpesvirus in Pacific oysters (*Crassostrea gigas*)

**DOI:** 10.1101/223032

**Authors:** Alejandro P. Gutierrez, Tim P. Bearf, Chantelle Hooped, Craig A. Stentort, Matthew B. Sanders, Richard K. Paley, Pasi Rastas, Michaela Bryrom, Oswald Matika, Ross D. Houston

## Abstract

Ostreid herpesvirus (OsHV) can cause mass mortality events in Pacific oyster aquaculture. While various factors impact on the severity of outbreaks, it is clear that genetic resistance of the host is an important determinant of mortality levels. This raises the possibility of selective breeding strategies to improve the genetic resistance of farmed oyster stocks, thereby contributing to disease control. Traditional selective breeding can be augmented by use of genetic markers, either via marker-assisted or genomic selection. The aim of the current study was to investigate the genetic architecture of resistance to OsHV in Pacific oyster, to identify genomic regions containing putative resistance genes, and to inform the use of genomics to enhance efforts to breed for resistance. To achieve this, a population of ~1,000 juvenile oysters were experimentally challenged with a virulent form of OsHV, with samples taken from mortalities and survivors for genotyping and qPCR measurement of viral load. The samples were genotyped using a recently-developed SNP array, and the genotype data were used to reconstruct the pedigree. Using these pedigree and genotype data, the first high density linkage map was constructed for Pacific oyster, containing 20,353 SNPs mapped to the ten pairs of chromosomes. Genetic parameters for resistance to OsHV were estimated, indicating a significant but low heritability for the binary trait of survival and also for viral load measures (h2 0.12 – 0.25). A genome-wide association study highlighted a region of linkage group 6 containing a significant QTL affecting host resistance. These results are an important step towards identification of genes underlying resistance to OsHV in oyster, and a step towards applying genomic data to enhance selective breeding for disease resistance in oyster aquaculture.

## Introduction

A specific genotype of the ostreid herpesvirus (OsHV-1-μvar) has been suggested to be the main cause of periodic mass mortality events in farmed Pacific oysters *(Crassostrea gigas)* worldwide (Segarra *et al*. 2010), with other contributing factors potentially including *Vibrio* bacterial infection and elevated temperature (Petton *et al*. 2015; Malham *et al*. 2009). Given that Pacific oysters account for 98% of global oyster production, which was estimated at ~0.6M tons in 2015, this pathogen is a significant problem for global aquaculture. Due to the current lack of effective options to prevent or control disease outbreaks (e.g. no option for vaccination and limited evidence of effective biosecurity) improving host resistance to OsHV-1 via selective breeding has become a major target.

A significant additive genetic component has been described for survival during OsHV-1 infection, with estimated heritability values ranging from 0.21 to 0.63 (Azéma *et al*. 2017; Camara *et al*. 2017; Dégremont *et al*. 2015a). Substantial efforts are being made to establish selective breeding programs for *C. gigas* with OsHV-1 resistance as the primary target trait. An encouraging response to selection for resistance has been observed in *C. gigas* spat after four generations of mass selection (Dégremont *et al*. 2015b). Modern selective breeding programs for aquaculture species can facilitate the simultaneous selection of multiple traits, including those not possible to measure directly on selection candidates. Genomic tools can facilitate this process, allowing for increase in selection accuracy and rate of genetic gain for target traits, with improved control of inbreeding (Houston 2017). Further, these tools allow investigation of the genetic architecture of key production traits, opening up possibilities for downstream functional studies to discover genes contributing directly to genetic variation. Putative QTL affecting host resistance to OsHV-1 have been identified using a linkage mapping approach (Sauvage *et al*. 2010), but genome-wide association approaches have not previously been performed in oysters and offer a substantially higher marker density and mapping resolution.

SNP arrays are enabling tools for genetic analysis and improvement of complex traits in farmed animal species. In the past few years, many genomic resources have been developed for *C. gigas* and include a reference genome assembly (Zhang *et al*. 2012), and a moderate number of genetic markers, such as microsatellites (Li *et al*. 2003; Sekino *et al*. 2003; Sauvage *et al*. 2009) and SNPs (Fleury *et al*. 2009; Sauvage *et al*. 2007; Wang *et al*. 2015). Additionally, low to medium density linkage maps have been developed, containing both microsatellites and SNPs (Li and Guo 2004; Sauvage *et al*. 2010; Hedgecock *et al*. 2015; Hubert and Hedgecock 2004). Importantly, the recent development of medium and high density SNP arrays for oysters (Gutierrez *et al*. 2017; Qi *et al*. 2017) raises the possibility of rapidly collecting genotype data for many thousands of SNP markers dispersed throughout the genome. These tools therefore facilitate development of high density linkage maps and high resolution genome-wide association studies into the genetic architecture of traits of economic interest. In addition, such genome-wide genotyping platforms enable testing of genomic selection approaches which are increasingly common in aquaculture breeding, with encouraging empirical data supporting the advantage over pedigree-based approaches (Tsai *et al*. 2015; Tsai *et al*. 2016; Vallejo *et al*. 2017; Dou *et al*. 2016; Correa *et al*. 2017).

The aim of this study was to investigate the genetic architecture of resistance to OsHV-1 infection in *C. gigas* using a large immersion challenge experiment followed by a GWAS to identify loci associated with the trait, and the relative contribution of these loci to the heritability of the trait.

## Methods

### Source of oysters and disease challenge

Oysters used in this study were obtained from multiple crosses of parents provided by Guernsey Sea Farms (UK) and reared at Cefas facilities. These comprised three pair crosses that were created at Cefas (from 3 sires and 2 dams) and each reared separately, while the rest of the crosses (from 14 sires and 14 dams) were obtained as spat from Guernsey Sea Farms and combined into a mixed culture tank at Cefas. Oysters were held at 20 +/− 2 C during post-settlement and fed with a combination of *Isocrysis, Tetraselmis, Chaetoceros, Pavlova sp*., and ‘Shellfish Diet 1800’ until they reached an appropriate size at approximately eight months of age. A subsample of approximately 1,000 oysters were then transferred to a new tank at 20 +/− 2 °C for two days for acclimation. An aliquot of the oyster herpes virus OsHV-1 μvar (amplified by two passages through *C. gigas*, purified by filtration and whole genome sequenced for confirmation using Illumina MiSeq technology) was then added to the water tank at an end concentration of 2.49x10^7^ copies /ml (empirically assessed by qPCR) with continuous flow. The challenge lasted for 21 days, by which time mortality rate had returned to baseline levels, and mortalities and survivors were snap-frozen and stored for DNA extraction.

### Phenotypic measurements

Survival was coded as binary trait i.e. 0 (mortality) or 1 (survival). The viral count of all samples was determined by qPCR according to (Martenot *et al*. 2010), with the addition of a plasmid based standard curve cloned for absolute quantification. The estimated copy number was then divided by the weight of the animal (mg) to obtain a measure of the viral load. Viral load values were then normalised by transformation to the logarithmic scale for further analyses.

### SNP array genotyping

Genomic DNA was extracted from the whole oyster (minus the shell) using the RealPure genomic DNA extraction kit (Valencia, Spain), quantified on Qubit and the DNA integrity was checked on a 1% agarose gel. Genotyping was carried out at Edinburgh Genomics using the recently developed Affymetrix SNP array for oysters (Gutierrez *et al*. 2017). After considering available DNA quality and quantity, only 897 samples were retained for genotyping (33 parents + 864 challenged offspring). After quality control (QC) using the Axiom Analysis Suite v2.0.0.35, 854 samples were retained following the “best practices workflow” with a sample and SNP call threshold of 90 resulted in 23,388 SNPS classified as good quality (‘PolyHighResolution’ and ‘NoMinorHom’ categories), from ~40K putative available for *C.gigas* on the array, and retained for downstream analyses.

### Linkage mapping

Linkage maps were constructed using Lep-map 3 (Rastas 2017). Families used for the generation of this map were assigned using Cervus (Kalinowski *et al*. 2007) as described by Gutierrez *et al*. (2017), and further confirmed through the IBD module in Lep-map3. Putative erroneous or missing parental genotypes were re-called using the “ParentCall2” module. Linkage groups were identified using the “SeparateChromosomes2” module using a LodLimit=60 and distortionLod=1. Data were then filtered to remove markers from families showing deviations expected Mendelian segregation ratios (“dataTolerance=0.001”) and used with the “OrderMarkers2” module to order the markers in the linkage groups. Individuals showing excessive recombination were also removed from the data as this indicated a potential problem with genotyping or family assignment for this individual. The estimated genome coverage of the map was calculated as c = 1 - e^−2dn/L^, where d is the average spacing of markers, n is the number of markers, and L is the length of the linkage map (Bishop *et al*. 1983). Only full sibling families were used for the construction of the linkage maps.

### Model and heritability estimation

Genetic parameters for the resistance traits were estimated using a linear mixed model approach fitting animal as a random effect using ASReml 4 (Gilmour et al. 2014) with the following model:

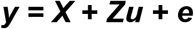

where **y** is the observed trait, **u** is the vector of additive genetic effects, **e** is the residual error, and **X** and **Z** the corresponding incidence matrices for fixed effects and additive effects, respectively. The (co)variance structure for the genetic effect was calculated either using pedigree **(A)** or genomic **(G)** matrices (i.e. **u** ~ N(0, Aσ_a_^2^) or N(0, Gσ_a_^2^)), where G is the genomic kinship matrix and σ^2^ is the genetic variance. Hence, the narrow sense heritability was estimated by the additive genetic variance and total phenotypic variance, as follows:

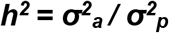

where σ^2^_a_ is the additive genetic variance and σ^2^_p_ is the total phenotypic variance which is a sum of σ^2^_a_ + σ^2^_*e*_. Heritability on the observed binary scale obtained for survival was converted to the underlying liability scale according to Dempster and Lerner (1950). The genomic relationship matrix required for the analysis was obtained according to (VanRaden 2008) using the GenABEL package (Aulchenko *et al*. 2007) and inverted using a standard ‘R’ function.

### Genome-wide association studies

The GWAS was performed using the GenABEL package (Aulchenko *et al*. 2007) in R. The genotype data were filtered as part of quality control by using the *check.markers* module to retain SNPs with a MAF > 0.01, call rate >0.90 and allow a deviation from Hardy-Weinberg Equilibrium < 1 x 10^−5^, leaving 16,223 filtered SNPs for downstream analyses. Association analyses were run using the family-based score test for association (FASTA) using the mmscore function (Chen and Abecasis 2007) with the mixed linear model (MLM) approach used to avoid potential false positive associations derived from population structure. Genotype data were used to calculate the genomic kinship matrix which was fitted in the model alongside the top four principal components as covariates to account for population structure. Additionally, the GWAS was run using the Efficient Mixed-Model Association eXpedited (EMMAX) software (Kang *et al*. 2010) to perform a form of validation test for SNPs identified as significant in the GenABEL analysis. The genome-wide significance threshold was set to 3.08x10^−6^ as determined by Bonferroni correction (0.05 / N), where N represents the number of QC-filtered SNPs across the genome, while the suggestive threshold was set as 3.08 x 10^−5^ (0.5/N), i.e. allowing 0.5 false positive per genome scan.

### Identification of candidate genes

To identify candidate genes potentially underlying the identified QTL for further study, the location of the most significant SNPs on individual contigs and scaffolds was recorded on the *C. gigas* genome v9 assembly (GCA_000297895.1) (Zhang *et al*. 2012). The sequences of these scaffolds / contigs were then aligned (using a custom-built blastn database) with the *C. gigas* gene annotation database. Contig and scaffold sequences for significant SNPs were also aligned using blastn and blastx (using non-redundant protein sequences) from the NCBI database.

### Data availability

Linkage map including all mapped markers and their position is given in File S1. Genotype data corresponding to all informative markers for all the individuals involved in this study is given in File S5.

## Results

### Challenge outcome and trait heritability

At the end of the 21 day disease challenge, 749 oysters had survived while 251 had died during the experiment. From the latter, 71 oysters had no body tissue at the moment of their removal, leaving 181 mortalities suitable for downstream analyses. Therefore, overall mortality was approximately 25 %, but in the subset of oysters available for genotyping the mortality was ~18%.

A total of 23 full sibling families were identified using the family assignment software. The largest comprised 231 individuals, and the smallest only two individuals. The vast majority of offspring were assigned to a unique parent pair, but a total of seven individuals were assigned to only one parent (five only to a dam and two only to a sire). Making use of the pedigree information, the estimated heritability on observed scale was 0.13 (0.06), corresponding to a value of 0.25 on the underlying liability scale (Table 1). These estimates were slightly lower when using the genomic kinship matrix, with 0.08 ± 0.03 and 0.17 for the observed and liability scale respectively. For viral load, heritability based on pedigree was estimated at 0.19 ± 0.08 and 0.13 ± 0.05 for genomic matrix. (Table 1).

**Table 1.**
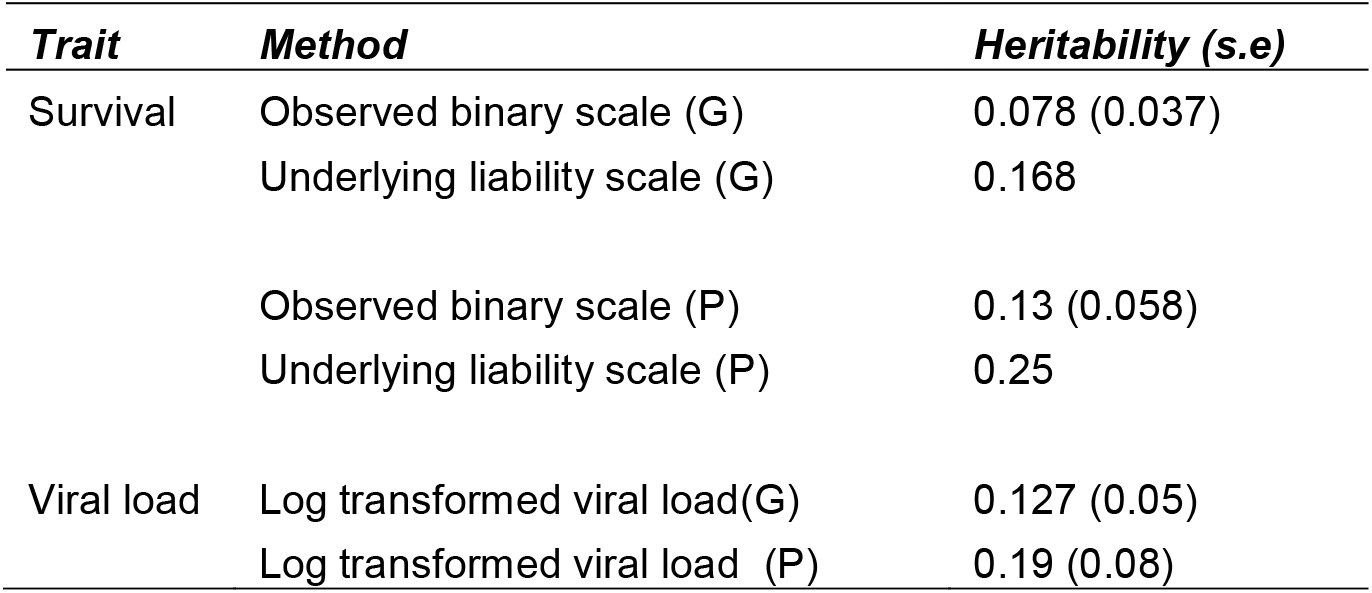
Estimated heritabilities for survival and viral load in challenged populations.

### Linkage map

The linkage mapping was performed using the 23 full sibling families comprising 809 progenies and 31 parents. On average 10% of the markers showed evidence of segregation distortion (p < 0.001) in at least one family with at least ten progenies, leaving 21,087 maternally informative markers and 20,528 paternally informative markers for map construction (Table S1).

The linkage map contains 20,353 SNPs distributed on 10 LGs (in accordance with the *C. gigas* karyotype) as shown in Figure 1, with a length of 951 cM for the male map and 994 cM for the female map. The ~20K mapped SNPs correspond to 1,921 scaffolds and 149 contigs, according to the latest oyster genome assembly (GCA_000297895.1, Zhang *et al*. 2012, File S1). These scaffolds and contigs containing mapped SNPs covered approximately 87% of the reference genome length.

**Figure 1.**
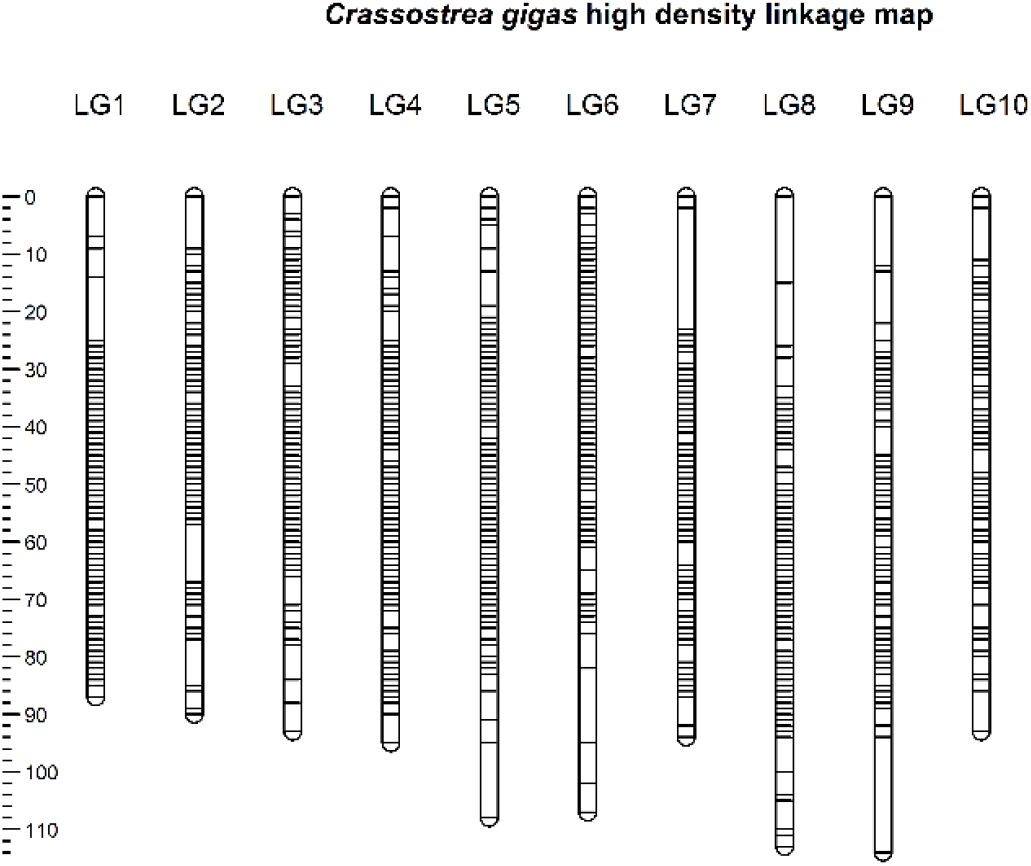
Distribution of SNP markers on the linkage map.

Linkage groups were labelled according to Hedgecock *et al*. (2015) to keep consistency across *C.gigas* linkage maps. Our medium density oyster array contains 464 of the SNPs mapped by Hedgecock *et al*. (2015). From these, 307 were mapped in the current study and their new linkage group assignment fully agrees with their previous assignment (File S2). Likewise, we observed that approximately 38 % (734 out 1,921) of the scaffolds with informative markers show evidence of errors in the assembly, due to assignment to at least two distinct LGs in our map (File S3). As expected, the number of LGs associated with scaffolds was positively correlated with scaffold length (Figure S1).

### Association analyses

The GWAS for the binary survival trait using the FASTA approach identified two markers showing a genome-wide significant association with the trait (both also significant using EMMAX, with an additional two SNPs significant using EMMAX only), as shown in Table 2, Figure 2 and File S4. Of the ten markers showing the most significant association in the two approaches, four markers are linked to LG 6 but they do not map to the same scaffold, nor are they close together on the linkage map. The proportion of phenotypic variation explained by the top ten markers ranged between 0.019 and 0.047, which implies a polygenic architecture to host resistance, albeit the LG 6 QTL potentially explains a large proportion of the genetic variance given the low heritability estimates.

**Figure 2.**
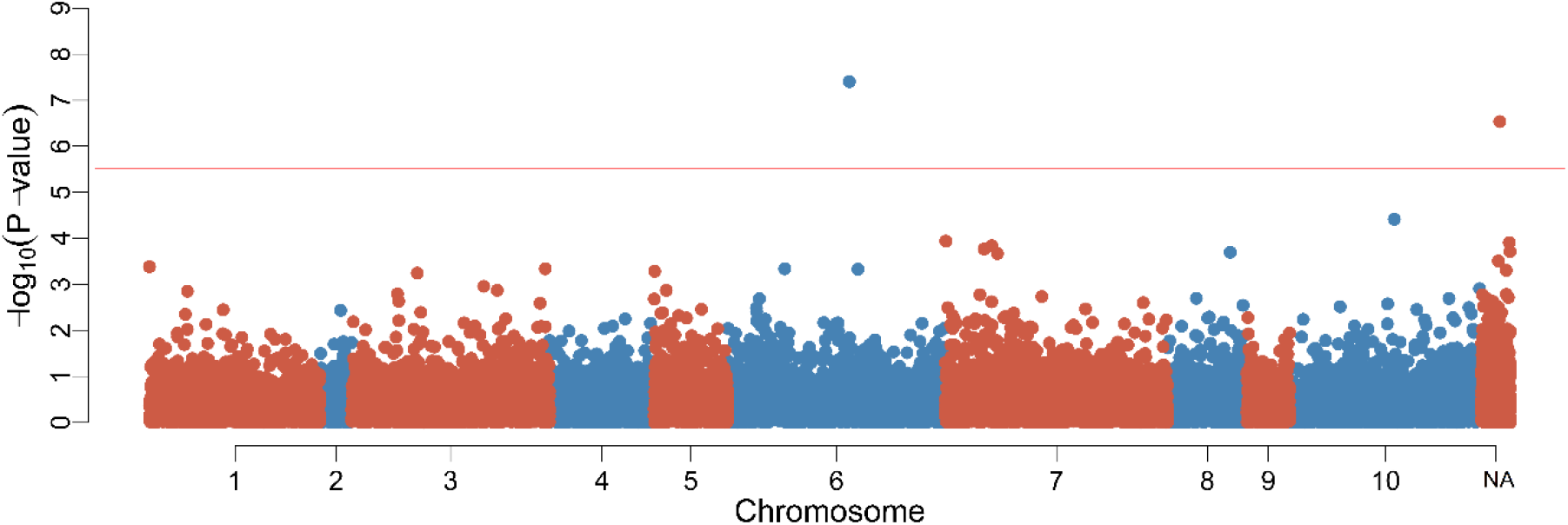
Manhattan plot the GWAS for survival. The position of the SNPs on the X axis is calculated according to the linkage map. “NA” represent a chromosome that contains markers not assigned to any linkage group. Horizontal red line indicates the genome-wide significance threshold.

**Table 2.** The top ten markers associated with survival.

| SNP ID | LG (position cM) | LG nearest marker (position cM) | Scaffold (position bp) | A1 | A2 | GenABEL | EMMAX | PVE | Nearest Gene |
| --- | --- | --- | --- | --- | --- | --- | --- | --- | --- |
| AX-169184215 | LG 6 (42.46) | - | scaffold241 (824,662) | T | G | 3.94E-08* | 4.74E-10* | 0.0473 | CORO1B |
| AX-169192574 | Unassigned | LG 6 (54.61) | scaffold1827 (350,776) | A | G | 2.91E-07* | 7.79E-08* | 0.0411 | MYO10 |
| AX-169208860 | Unassigned | LG 1 (54.37) | scaffold714 (58,763) | G | A | 0.000124 | 5.72E-07* | 0.0224 | CYP1A1 |
| AX-169209993 | LG 7 (9.48) | - | scaffold1599 (493,016) | T | C | 0.000115 | 1.56E-06* | 0.0231 | D2R |
| AX-169207075 | LG 5 (47.54) | - | scaffold57 (142,065) | C | T | 0.004125 | 1.09E-05 | 0.0122 | IFT88 |
| AX-169210119 | Unassigned | LG 6 (29.41) | scaffold198 (583,825) | T | C | 0.000194 | 2.25E-05 | 0.0206 | RANBPM |
| AX-165319118 | LG 5 (25.77) | - | scaffold43494 (138,038) | G | A | 0.000519 | 4.82E-05 | 0.019 | KPNA1 |
| AX-169158711 | LG 6 (42.59) | - | scaffold109 (558,765) | G | A | 0.000468 | 6.48E-05 | 0.0183 | CASP |
| AX-169199571 | LG 10 (42.24) | - | scaffold186 (320,367) | C | T | 3.85E-05 | 7.02E-05 | 0.0247 | AP1AR |
| AX-169168346 | LG 3 (43.83) | - | scaffold1785 (251356) | G | T | 0.000568 | 8.37E-05 | 0.018 | KIF6 |

\* Genome-wide significant ( $p < 0.05$ ) markers. A1 & A2, major and minor allele. PVE, phenotypic variation explained by the SNP. The physical position of the SNPs on the Scaffolds are given according to the Pacific oyster reference assembly (Genbank ID GCA\_000297895.1).

The GWAS for the trait of viral load detected two markers showing significant genome-wide association with both FASTA and EMMAX, with an addition eight SNPs identified as significant using EMMAX only (Table 3, Figure S2 and File S4). The SNP showing the most significant association is located in LG 8, however, no other markers are located in the same LG. While most of the markers significantly associated with the trait were not mapped, the nearest mapped SNPs according to their position on the genome scaffolds suggests that three SNPs are located on LG 6. Therefore, it is plausible that there is at least one QTL on LG 6, and this QTL may affect both viral load and the binary trait of survival. The proportion of phenotypic variation in viral load explained by the top ten markers ranged between 0.0209 and 0.037.

**Table 3.** Top ten markers associated with viral load.

| SNP ID | LG (position cM) | LG nearest marker (position cM) | Scaffold (position bp) | A1 | A2 | GenABEL | EMMAX | PVE | Gene |
| --- | --- | --- | --- | --- | --- | --- | --- | --- | --- |
| AX-169203956 | LG 8 (0) | - | scaffold501 (742,989) | C | T | 6.54E-07* | 3.47E-09* | 0.037 | FBN2 |
| AX-169210119 | Unassigned | LG 6 (29.41) | scaffold198 (583,825) | T | C | 9.09E-07* | 9.33E-08* | 0.0349 | RANBPM |
| AX-169172429 | Unassigned | LG 4 (57.01) | scaffold713 (269,794) | T | G | 3.72E-06 | 9.36E-07* | 0.0314 | B3GALNT2 |
| AX-169192982 | Unassigned | LG 6 (43.46) | scaffold1093 (208,087) | C | T | 9.61E-06 | 1.54E-07* | 0.0284 | SKI |
| AX-169167580 | Unassigned | LG 6 (36.95) | scaffold1763 (82,048) | A | G | 1.65E-05 | 8.91E-07* | 0.0277 | CLEC16A |
| AX-169199878 | Unassigned | LG 8 (71.91) | scaffold536 (135,453) | T | C | 2.11E-05 | 1.42E-06* | 0.0269 | TNR |
| AX-169203386 | Unassigned | LG 10 (52.04) | scaffold1301 (188,626) | C | T | 2.27E-05 | 1.13E-06* | 0.0266 | RAPGEF2 |
| AX-169196070 | Unassigned | LG 1 (57.32) | scaffold433 (1,082,890) | G | A | 3.62E-05 | 1.69E-06* | 0.0261 | CARS |
| AX-169199276 | LG 7 (6.68) | - | scaffold128 (550,765) | G | T | 3.70E-05 | 8.44E-07* | 0.0243 | SMARCA5 |
| AX-169194053 | LG 4 (19.39) | - | scaffold728 (174,857) | G | T | 0.000148 | 2.22E-06* | 0.0209 | TNIK |
| AX-169193982 | LG 1 (47.02) | - | scaffold41452 (35,018) | A | G | 6.79E-05 | 6.03E-06 | 0.0243 | U/P<sup>a</sup> |
| AX-169192459 | Unassigned | LG 1 (22.66) | scaffold447 (373,487) | G | A | 7.28E-05 | 9.00E-06 | 0.0232 | SCARF2 |

\* Genome-wide significant ( $p < 0.05$ ) markers. A1 & A2, major and minor allele. PVE, phenotypic variation explained by the SNP. <sup>a</sup> U/P indicates uncharacterized protein. The physical position of the SNPs on the Scaffolds are given according to the Pacific oyster reference assembly (Genbank ID GCA\_000297895.1).

## Discussion

### Heritability of OsHV-1 resistance

Estimates of heritability observed for survival to OsHV-1 challenge in the current study were low to moderate (0.078-0.25) in comparison to other recent studies that have analysed resistance to OsHV-1, where estimates have ranged from 0.21 to 0.63 (Dégremont *et al*. 2015a; Azéma *et al*. 2017; Camara *et al*. 2017). Mortality resulting from OsHV-1 exposure in our challenge was relatively low, reaching ~ 25% in the overall challenge. The mortality level in the genotyped samples was lower (~18%), although it is not clear if the dead oysters found with no tissue were affected by the virus or were abnormal at the time of the exposure. It is possible that the population studied may have high level of innate resistance to OsHV-1, considering the low mortality level in ~8 month old oysters compared to the mortalities typically observed due to OsHV-1 exposure in spat and juvenile oysters (Azéma *et al*. 2017). Oysters from these families also showed lower mortality levels compared to other batches of oyster spat when using a more established single animal bath OsHV-1 challenges (data not shown), which would support the possibility of a relatively resistant sample of animals.

### Linkage map

The linkage map construction resulted in 10 linkage groups that correspond to the number of chromosomes of *C. gigas*, successfully mapping ~20K SNPs. The highest density linkage map for *C. gigas* to date was described by Hedgecock *et al*. (2015) and contains ~1.1K SNPs and microsatellites. Therefore, the linkage map presented in the current study is an improvement to existing resources offering an advance for oyster genomics with potential in assisting future mapping studies, particularly those using the medium density SNP array.

Family assignments were rigorously tested to avoid pedigree errors in the construction of the linkage maps. Distortions from the expected Mendelian segregation were found in ~10 % of the SNPs in the larger families (p < 0.001) (Table S1). Moderate levels of segregation distortion have been commonly observed in oysters (Jones *et al*. 2013; Hedgecock *et al*. 2015; Guo *et al*. 2012) and bivalves in general (Saavedra and Bachère 2006). In the current study, distorted markers were included for the linkage group assignment, but were filtered out for the determination of the order in the LG. It has been argued that distorted markers can affect marker ordering, albeit the effect on map construction has been shown to be minor (Hackett and Broadfoot 2003; Guo *et al*. 2012).

A measure of the quality of the linkage map was given by overlap with a previous linkage map described by Hedgecock *et al*. (2015). Several hundred SNPs were successfully re-mapped to the same LG, indicating correct LG definition.

Accordingly, assembly errors observed by Hedgecock *et al*. (2015) were also observed in our high-density linkage map, where almost ~40% of the mapped scaffolds were assigned to more than one LG (File S1). This linkage map should be able to provide a good base for the identification of assembly errors and the potential re-assembly of the genome, which seems like a requirement to maximise its utility for future genomics research in this species.

### GWAS and associated genes

The association analyses for OsHV-1 survival and viral load suggest that both traits are likely to be impacted by multiple genomic regions, albeit the putative QTL on LG 6 potentially explains a large proportion of the genetic variation. Accordingly, GWAS for survival found SNPs surpassing the genome-wide threshold on LG 6, and SNPs surpassing the suggestive threshold on LG 1, LG 5, & LG 7 (Figure 2, Table 2 and File S4). For the trait of viral load, markers showing a genome-wide significant association were located in LG 8, LG 6LG 10 & LG 4, and suggestive association found in LG 1 & LG 7(Table 3 and File S4). The only previously published study describing genomic regions associated to summer mortality resistance found significant QTL in LG V, VI, VII & IX (which correspond to LG 6, LG 7, LG 8 & LG 10 in our map) in different families (Sauvage *et al*. 2010). It is noteworthy that LG 6 contains genome-wide significant SNPs for both survival and viral load (and was previously detected by Sauvage *et al*. 2010). In addition, a single SNP (AX-169210119) reached genome-wide significant level for viral load, and the suggestive level for survival. While this SNP was not mapped directly, the nearest mapped SNP was linked to LG 6.

Numerous genes were identified from the genomic regions flanking the most significant SNPs impacting the resistance traits. While the limits defined for screening flanking regions of significant SNPs were defined practically (i.e. the contig / scaffold to which the SNP maps), these genes may represent candidates for future validation, resequencing and functional testing. The SNP showing an association with both survival and viral load (AX-169210119) was located in the RAN Binding Protein 9-like gene which has recently been linked to the interferon gamma signalling pathway (Zhang *et al*. 2017), and also in viral adhesion and its replication in host cells (Yang *et al*. 2015). Another gene located near a significant SNP (AX-169184215) is a Coronin gene (CORO1B), from a family of genes that have multifaceted roles in immune response (Tokarz-Deptuła *et al*. 2017). Finally, the actin motor protein Myo10 gene is located near AX-169192574, and this gene encodes a protein which is essential for release of Marburgvirus particles from host cells (Kolesnikova *et al*. 2007). These and other genes may form the basis for downstream functional studies to assess their function in host response to virus in oysters. In addition, validation studies are required in independent populations to assess the robustness of the observed association between the significant SNPs and OsHV-1 resistance in oysters. Further, from a practical breeding perspective, these SNPs may have potential for marker-assisted or genomic selection to improve host resistance in farmed oyster populations.

## Conclusion

This study is the first to report GWAS using the a high density SNP panel Pacific oysters, and was enabled by the recent development of a SNP array (Gutierrez *et al*. 2017). Heritability of resistance to OsHV-1 in oysters was significant, but low to moderate in magnitude. The fact that this heritability was detected using both the pedigree and genomic relationship matrix implies that selective breeding and genomic selection for resistance is likely to be effective. Using the genotype data, a high-density linkage map was constructed for *C. gigas*, and the GWAS identified numerous markers showing a genome-wide significant association with the traits.

The most encouraging QTL was located on LG 6, reaching genome-wide significance for the binary trait of survival, with some evidence of a significant association with viral load. Future analyses will test candidate genes identified by the GWAS, verify trait-associated SNPs in independent populations, and test genomic selection as a tool to enhance host resistance to this problematic pathogen for oyster aquaculture.

## Acknowledgements

The authors gratefully acknowledge funding from BBSRC and NERC under the UK Aquaculture Initiative (BB/M026140/1, NE/P010695/1) in addition to BBSRC Institute Strategic Programme Grants (BB/P013759/1 and BB/P013740/1). SNP array genotyping was carried out by Edinburgh Genomics, The University of Edinburgh. Edinburgh Genomics is partly supported through core grants from NERC (R8/H10/56), MRC (MR/K001744/1) and BBSRC (BB/J004243/1).

